# Towards understanding the structure of the capsid of Banana Bunchy Top Virus

**DOI:** 10.1101/2020.02.12.945212

**Authors:** Sundaram Sairam, Ramasamy Selvarajan, Savithri S Handanahalli, Sangita Venkataraman

## Abstract

Banana is the major staple food crop for approximately 400 million people. Bunchy Top disease of Banana is one of the most devastating diseases caused by Banana Bunchy Top Virus (BBTV) that results in a significant loss of yield, stunting and bunchy appearance of leaves. While many isolates of BBTV from various regions of India have been characterized by different groups, no structural study exists for this important virus. To pursue structural studies, the pET28a clone of coat protein (CP) gene from BBTV isolate of Hill Banana grown in lower Pulney Hills (Virupakshi) of Tamilnadu was expressed in BL21 (DE3) pLysS. Purification of the CP was done using Ni-NTA affinity chromatography. In vitro capsid assembly studied using sucrose density gradient centrifugation suggested that the CP did not assemble as virus like particle (VLPs) but remained as smaller oligomers. Studies using dynamic light scattering (DLS) indicates that the purified protein is poly-dispersed represented majorly as pentamers. Studies using both homology modelling and *ab initio* structure determination gave useful insights into the probable fold of the CP suggesting it is a β-sandwich fold similar to that seen in majority of plant viruses. *In silico* capsid reconstruction aided understanding of the quaternary organization of subunits in the capsid and molecular interactions present between the subunits. The location of aphid binding EAG motif was identified on the surface loops close to the pentameric axis indicating their role in vector mediated transmission.

## Introduction

X-ray diffraction studies on single crystals of viruses enable visualization of the structures of intact virus particles at near-atomic resolution. These studies provide detailed information regarding the CP folding, capsid architecture, molecular interactions between protein subunits, and plausible sites of receptor recognition (Prasad & Schmid, 2012; Venkataraman et al., 2008). Such learning is pivotal in designing strategies for combating infections due to viruses in plants, animals and humans. Banana and plantains are grown in about 120 countries in mixed cropping systems by small holders and occasionally in monoculture (INIBAP, 2002). India is the largest producer of Banana in the world. Of 1919 million tonnes of banana that was produced globally in 2018, India accounted for 116.1 million tonnes (*FAO. 2019. Banana Statistical Compendium 2018. Rome.*, n.d.). Banana is an important fruit crop of many tropical and subtropical regions of the world including India, Philippines and China. BBT disease of Banana is one of the most devastating diseases caused by BBTV that results in a significant loss of yield, stunting and bunchy appearance of leaves (Figure 1) (Dale, 1987; Qazi, 2016; Vetten, 2008).

**Figure 1:**
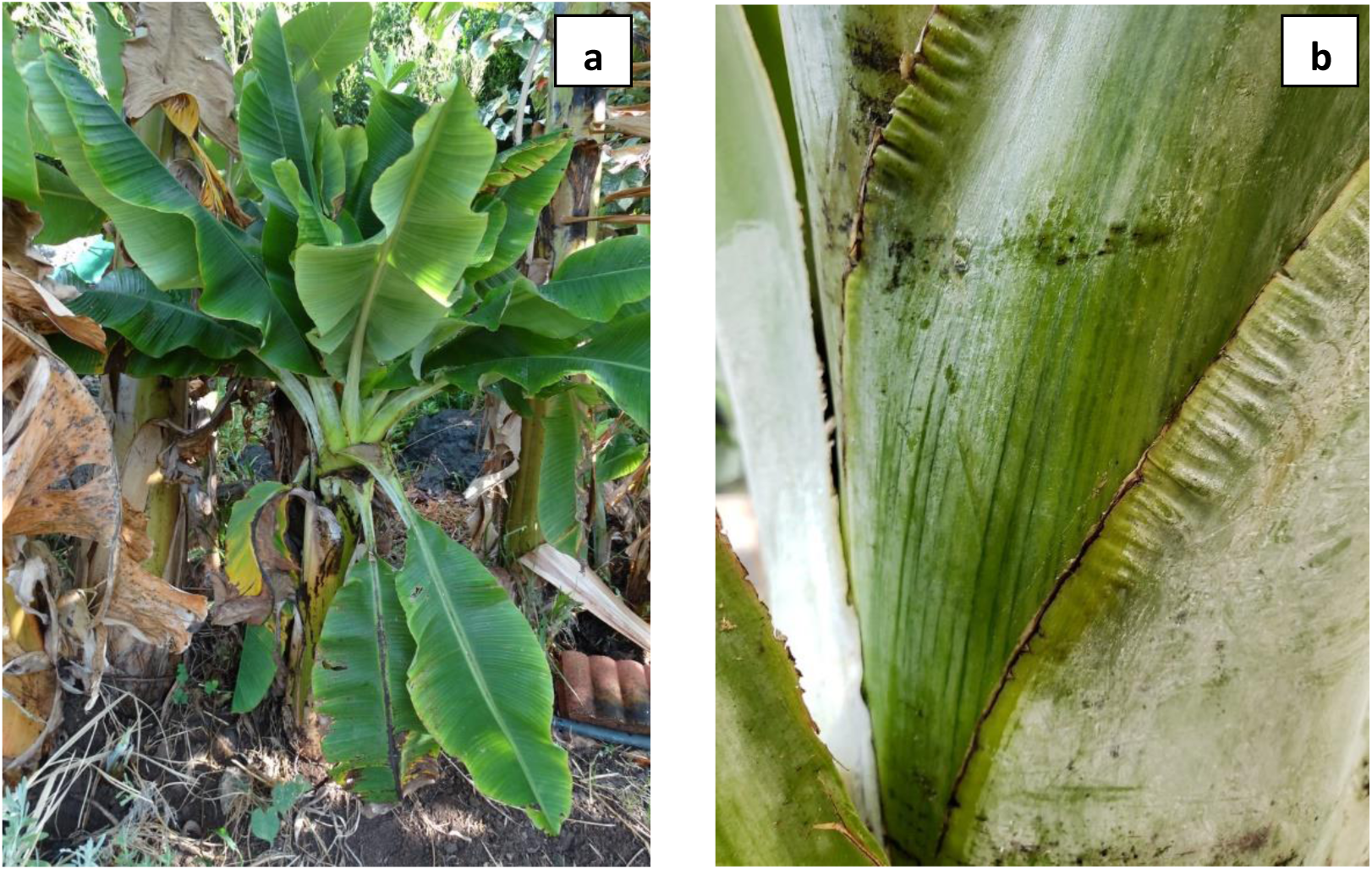
Bunchy top disease of banana caused by BBTV. a) Banana plant showing stunted bunchy appearance due to BBTV infection (https://www.flickr.com/photos/scotnelson/42839451145). b) Infected leaf showing dark green dash pattern due to BBTV infection (https://www.flickr.com/photos/scotnelson/28824232288).

BBTV is a member of *Nanoviridae* and represents the genus Babusvirus (Burns et al., 1995; Malathi & Renuka Devi, 2019). It belongs to group II viruses under Baltimore classification and is characterized by single stranded DNA (ssDNA) genome. The genome is multipartite consisting of six to eight circular segments that are separately encapsidated inside individual icosahedral shells of T=1 symmetry. The non-enveloped shell is a small round capsid of about 18 to 19nm in diameter and is made of viral CP, which is encoded by one of the segments of the genome, DNA-S. The virus is known to infect all members of Musaceae family and is transmitted by aphids ((King et al., 2012; Mandal, 2010; Qazi, 2016; Venkataraman & Selvarajan, 2019).

Globally, many groups have worked on characterizing BBTV from various countries like Pakistan (Amin et al., 2008), Africa (Kumar et al., 2011), China (Yu et al., 2011), Indonesia (Chiaki et al., 2015; Stainton et al., 2015), India (Banerjee et al., 2014; Selvarajan et al., 2010; Selvarajan & Balasubramanian, 2013; Vishnoi et al., 2009) and Srilanka (Wickramaarachchi et al., 2016). Since, BBTV is naturally transmitted by aphid vectors, a number of studies have been undertaken to get a deeper understanding of the mechanism of transmission (Hooks et al., 2009; Watanabe et al., 2016). There have been many research articles on the techniques of identification of BBTV in the fields. These include Direct Antigen Coated (DAC) ELISA and Double Antibody Sandwich (DAS) ELISA (Chen & Hu, 2013; Mansoor et al., 2005). Other sophisticated ways of detection involves tissues disruption from banana plants followed by real-time TaqMan(®) PCR that is highly sensitive and could detect as few as 2.76 copies of BBTV genomic DNA or 1.0 ng-1.0mg of infected banana leaves (Chen & Hu, 2013). Recently, paper-based gene based sensors have been developed for identifying contagious plant viruses in the fields (Wei et al., 2014). Since BBTV has a multipartite genome, different studies have focused on the recombination and reassortment ability of BBTV genome (Stainton et al., 2015). Fu *et al.*, (2009) have researched on the evolution of BBTV by studying the various recombination events (Fu et al., 2009). A recent study involving identification and *in silico* characterization of defective molecules associated with isolates of BBTV has suggested the involvement of defective DNA-R molecules (Stainton et al., 2015). An interesting study by Zhuang *et al.* has revealed the antagonistic effect of BBTV multifunctional protein B4 against Fusarium oxysporium (Zhuang et al., 2016). In a different study, Selvarajan and Balasubramaniam (Selvarajan & Balasubramanian, 2014) have cloned and sequenced the complete genome comprised of six DNA components of BBTV infecting Hill Banana grown in lower Pulney hills of Tamil Nadu State in India. Transgenic banana plant expressing siRNA to develop resistant varieties were developed by various groups (Elayabalan et al., 2015; Shekhawat et al., 2012). The replication initiation protein of Nanovirids esp of Abaca bunchy top virus (ABTV) and BBTV have been recently reviewed (Venkataraman & Selvarajan, 2019).

Considering the impact of Banana in our economy and growth, significant research and studies have been undertaken to prevent and treat diseases of bacteria, virus and fungi that affect banana. However, structural studies are lacking for this important class of virus infecting Banana plantation all over the world. Hence, the present study focuses on getting insights into the structure of BBTV CP and capsid by undertaking cloning of the CP in pET28a vector and overexpressing in bacterial system to purify sufficient protein for crystallographic studies. However, due to very low yield of CP protein, crystallization attempts have been unsuccessful so far. Modeling studies and *Ab initio* structure prediction of CP followed by *in silico* reconstruction of entire BBTV capsid provided deeper understanding of assembly, quaternary organization and interactions of BBTV. The results obtained from various studies are discussed in the current paper.

## Methodology

### Cloning studies

The infected leaf samples of BBTV were collected from the farmer’s field near National Research Center for Banana and confirmed by direct antigen coating (DAC) ELISA utilizing polyclonal antiserum of BBTV. The total DNA was extracted from the infected leaves through CTAB method (Selvarajan et al., 2008). The 513 bp gene of CP was amplified through PCR using standard CP specific primers (forward 5’-ATG GCTAGGTATCCGAAGAAATCC-3’ and reverse 5’ TCAAACATGATATGTAATTCTGTTC −3’). The initial denaturation was for 5min at 94°C; followed by 30s denaturation at the same temperature, annealing for 45 secs at 53°C, and extension at 72°C for 30 cycles. The PCR products were analyzed in agarose gel. The CP gene was purified post PCR and was cloned into pET-28a (+) vector between XhoI and NcoI sites and confirmed by sequencing (AF148945.1:227-739).

### Expression and Purification

The pET28a plasmid with BBTV CP gene was transformed into BL21 (DE3) pLysS, plated on LB-agar. Single colony was inoculated into LB broth and left O/N for growth at 37°C. 1% primary culture was used for inoculating 500 ml of LB broth (Secondary culture). The secondary culture was induced at 0.6 OD with IPTG at a concentration of 1M. Post induction, the culture was kept at 37° for 3hrs for protein expression. The expression was studied at various induction temperature, IPTG concentrations and against different bacterial strains such as Rosetta and codonPlus. The cells were harvested by centrifugation for 10 mins at 8000rpm. The pellet was suspended in the Lysis Buffer containing 20mM Tris (pH8) and sonicated using VibraCell ultrasonic processor at 50% amplitude/ 10 sec pulse/ 20 sec rest/ 20 min in ice. The soluble and insoluble fractions were separated following cell lysis. For every bacterial strain used, the lysis buffer composition was varied, and the solubility was assessed. Further purification was carried out using affinity chromatography using Ni-NTA resin. Two rounds of washing using buffers with 20mM and 30mM Immidazole concentration was carried out prior to elution with 300mM Immidazole.. The elute was subjected to dialysis to remove the imidazole and concentrated to 1ml. The final purified protein was checked on the gel, the concentration was assessed by absorbance at 280nm. The 260/280 ratio was taken for assessing the presence of nucleic acid.

### Protein Characterization

#### Density Gradient Centrifugation

The clear supernatant post cell lysis was transferred into 32 ml ultra-centrifuge tubes and spun at 26,000 rpm for 3 hours at 4°C in AH629 rotor. The ultra-pellet was suspended in a small volume of 20mM Tris buffer (pH 7.5) through end-to-end rotation, layered onto tubes with 10 – 40 % sucrose density gradient and centrifuged at 26,000 rpm for 3 hours. Following the run, 1.5 ml fractions were collected from the bottom of the gradient. The presence of VLPs was assessed by studying the presence of light scattering zone, monitoring the absorbance of the fractions at 280 nm and SDS-PAGE analysis (He, 2011).

#### Dynamic Light Scattering (DLS)

Protein solution, buffer, 100% EtOH, and MQ H2O were filtered through 0.1 µm filters prior to the experiment. The concentration of protein was kept at ∼ 1 mg/ml. DLS experiments were performed using Viscotek 802 DLS in a quartz cuvette (volume, 12 μl) at 22 °C.

#### In silico structure predictions

The amino acid sequence of BBTV CP (Q65386) was retrieved from the SwissProt database (Bairoch, 2000) and used as target for *ab initio* structure prediction and homology modelling using various servers that are available online or using commercial packages such as Schrodinger (Schrödinger, 2018). Since the protein sequence identity for the BBTV CP with all the available proteins of PDB was very poor (< 20%), a proper homologous model is unavailable. *Ab initio* structure prediction was done using iTASSER (Iterative Threading ASSEmbly Refinement) (Yang & Zhang, 2015), Robetta (Kim et al., 2004; Ovchinnikov et al., 2018) and Quark (Xu & Zhang, 2012, 2013), while homology modeling was attempted using Schrodinger (**Schrödinger Release 2019-4**: Prime, Schrödinger, LLC, New York, NY, 2019), SwissModeler (Bienert et al., 2017), RaptorX (Källberg et al., 2012, 2014), Phyre (Kelley et al., 2015; Kelley & Sternberg, 2009) and MMM (Jeschke, 2018). The models were subjected to quality evaluation using QMean server (Benkert et al., 2009, 2011) and SAVES (Pontius et al., 1996) for assessing structure quality. The best scoring model was considered for further analysis. The disorder was assessed using the PrDOS (Ishida & Kinoshita, 2007), and the nucleic acid binding was assessed using bindUP server (Paz et al., 2016).

### Model Validation

The finalized model was energy minimized using the protein-preparation wizard of the Schrodinger suite to corrects problems such as missing hydrogen atoms, incomplete side chains and loops, ambiguous protonation states, and flipped residues (Schrödinger, 2018). Following minimization, the model was exported to the SAVES server Version 5.0 (Pontius et al., 1996) and their overall stereochemical quality, including backbone torsional angles was checked through the Ramachandran plot using the program PROCHECK (Laskowski et al., 2012).

### VIPERdb analysis

In silico capsid reconstruction was done using VIPER and the capsid features were analyzed using the VIPERdb server (Carrillo-Tripp et al., 2009; Ho et al., 2018). The inner and outer radii are calculated by the server after deriving the minimum and maximum distance calculated from the origin (0,0,0) going over all the atoms. The number of inter-residue contact between a pair of residues from two (different) subunits was evaluated by VIPERdb using the criteria that the distance between the center of mass of the side chain atoms is within the distance criteria as estimated on the basis of the structures available in PDB (Godzik et al., 1992). Subunit association energies were estimated by the server as the product of atomic buried surface areas and the solvation parameters (Eisenberg & Hill, 1989; Horton & Lewis, 1992). The program CHARMM was used to calculate the Buried surface areas applying a probe radius of 1.4Å (B. R. Brooks et al., 2009; Bernard R. Brooks et al., 1983).

## Results and Discussion

### Cloning and expression results

The total DNA was extracted from the BBTV infected leaf sample and CP gene was amplified using the standard primers through PCR (Figure 2a). The CP gene was cloned into pET28a vector between XhoI and NcoI sites following purification of the PCR product (Figure 2b). The clone was also confirmed using sequencing. After standardization of expression, a 2l culture was lysed with solution containing 25mM Tris, 100mM Nacl, 1% Triton X 100 and 10mM Immidazole and further purified using affinity chromatography. Figure 3a and 3b show the overexpression of CP protein and the purified band following affinity chromatography, respectively. A band of 18.9 KDa corresponding to the CP was observed in 0.8% SDS PAGE (Figure 3b). The 260/280 ratio of 0.8 indicated the absence of nucleic acid in the preparation. To check if the expressed CP assembled into VLPs, the lysate was subjected to sucrose density gradient centrifugation (Figure 4a). No light scattering zone was observed. The absorbance of the fractions were recorded and the samples were run on a 0.8% gel. No bands were observed in the gel and the absorbance was insignificant indicating that the expressed BBTV-CP did not assemble into T=1 capsid.

**Figure 2:**
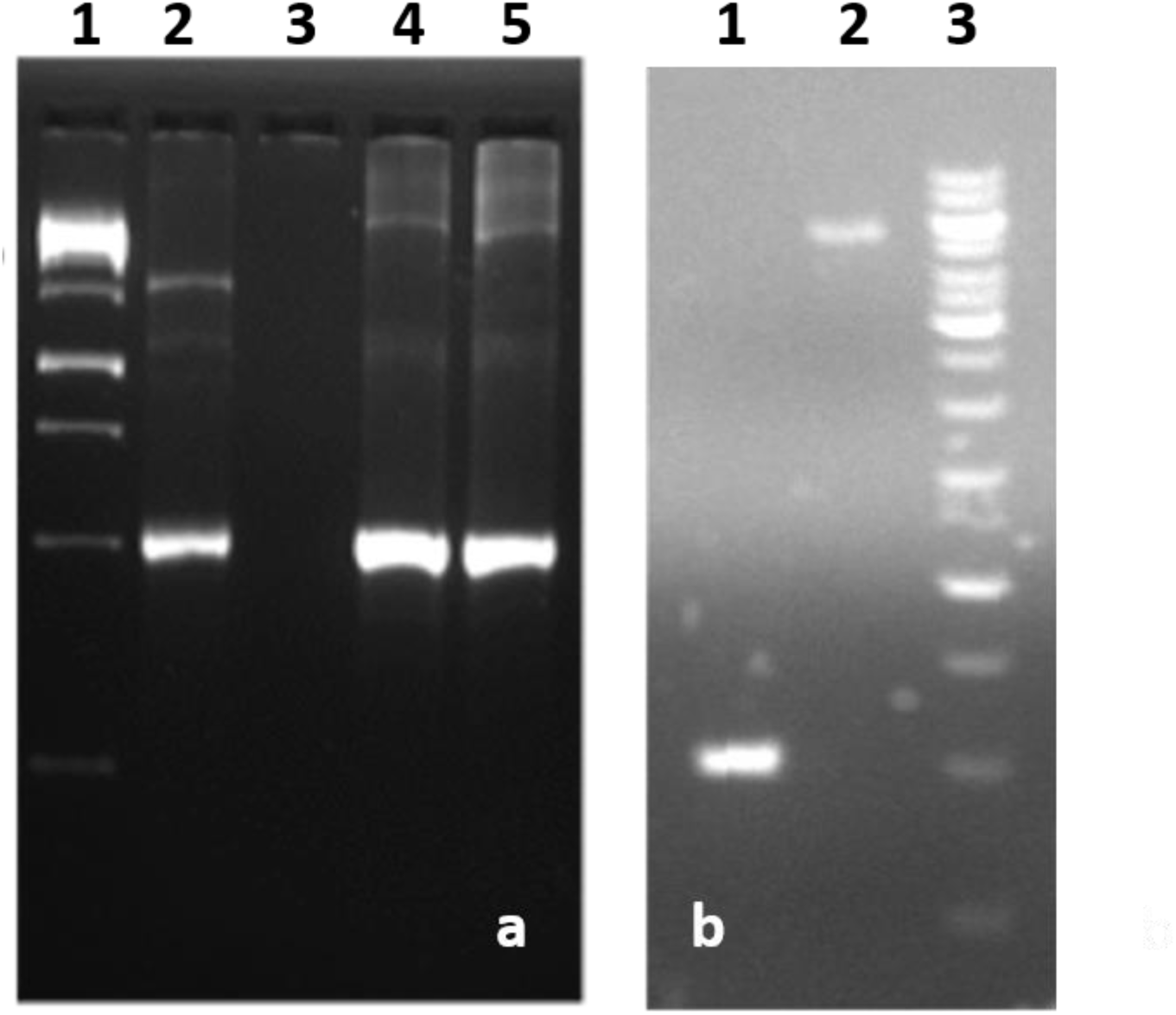
Cloning of CP gene in pET28a vector. a) The amplified CP gene of 513bp following PCR with DNA extracted from BBTV infected leaves. Lane 1 DNA marker, Lanes 2, 4 and 5 show the amplified CP band corresponding to 513bp. b) Confirmation of insert following cloning into pET28a. Lane 1 CP fragment released after restriction digestion with XhoI and NhoI, lane 2 Full length plasmid, lane 3 is the marker DNA.

**Figure 3:**
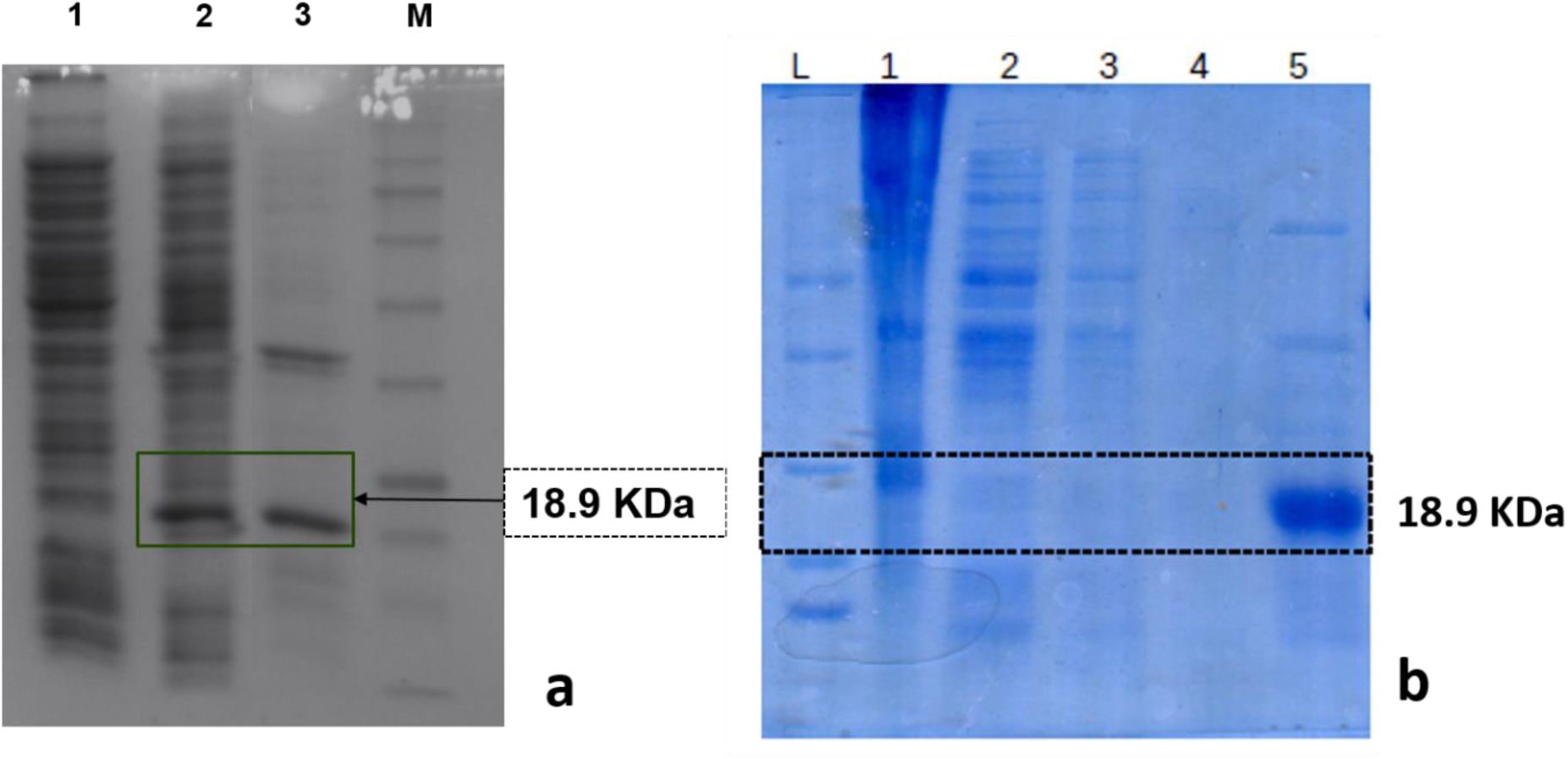
Expression and purification of CP gene in BL21 (DE3). a) Protein expression was induced by 1M IPTG kept at 37°C for 3 hours following induction. Lanes 1 shows uninduced sample, Lane 2 and 3 show the insoluble and soluble component following cell lysis. b) Lane 1 Protein marker, Lanes 2, 3, 4 and 5 wash through of Ni-NTA binding, Lane 6 eluted protein.

**Figure 4:**
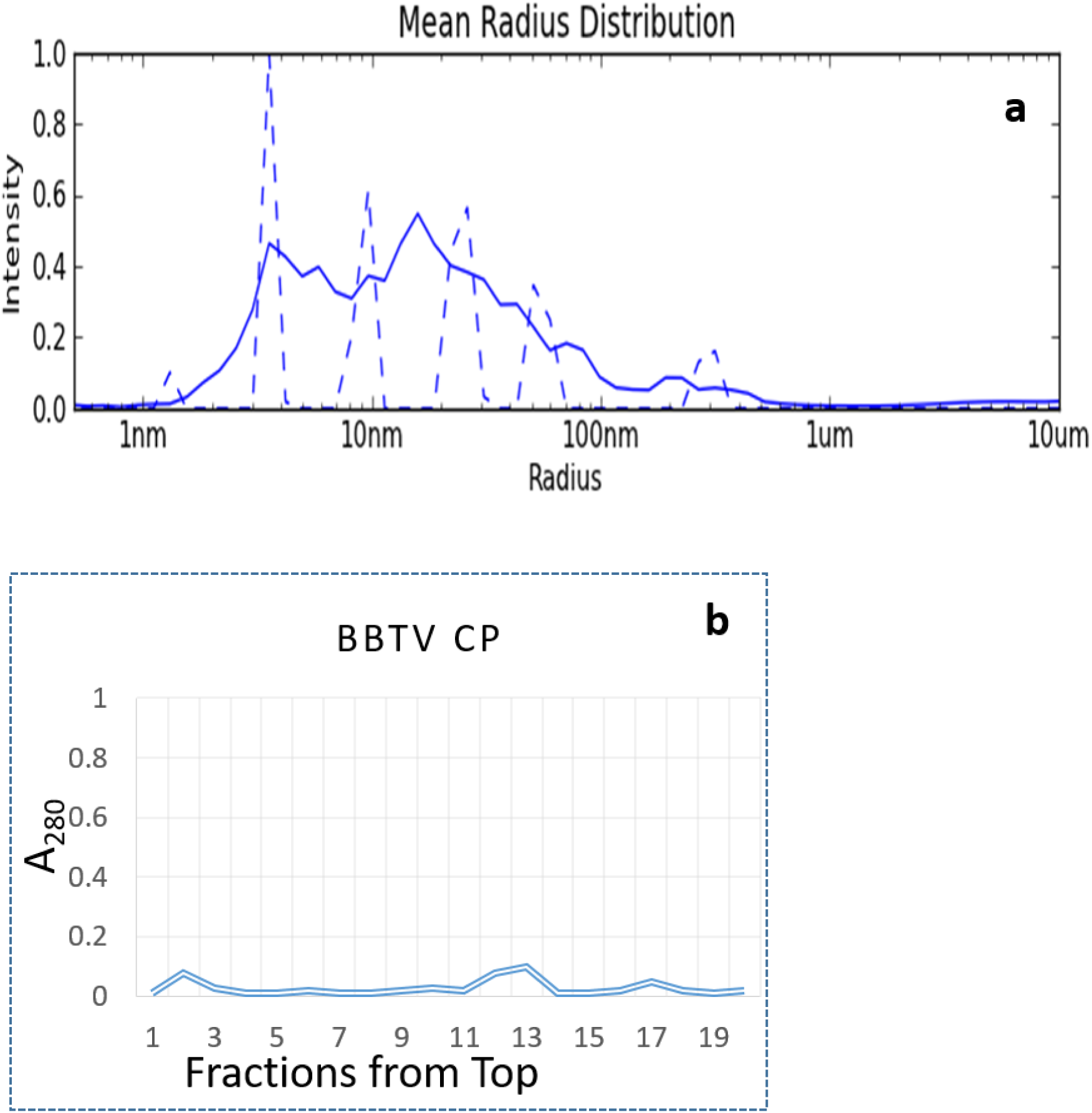
Protein homogeneity and assembly study a) DLS of purified CP showing the preponderance of pentamers. b) Sucrose density gradient profile showing the absence of VLPs in the protein preparation.

Unlike the native capsid, there was absence of any nucleic acid association indicating that specific interactions with ssDNA was probably mandatory for proper folding of the CP and eventual capsid assembly (Sangita et al., 2004; Satheshkumar et al., 2004). The DLS results also indicated that the CPs did not form regular capsids but instead were poly-dispersed with a greater population of pentamers over other species (Figure 4b). As the yield of the protein was insufficient for crystallization trials *in silico* methods were employed for gaining understanding of the nature and structure of the CP.

#### In silico modeling

Table 1 shows the results of *in silico* modeling obtained using both *ab initio* structure prediction and homology modeling with various servers. While the servers and programs predicted a consistent β structure for the overall model with regions of disorder at the amino terminus, there was not much of agreement between the other regions of the protein. While the coverage and QMean score was better for the *ab initio* model predicted by Robetta (Das & Baker, 2008; Ovchinnikov et al., 2018), they were less encouraging for others. Hence, the model predicted by Robetta was used for further analysis. The Robetta server used c-terminal domain of polymerase basic protein 2 from influenza virus (PDB ID: h5n1) as reference protein for building the model. The same protein was picked by ITASSER, Schrodinger and Phyre as the reference protein for modeling the structure.

**Table 1:** Comparison of the QMean scores of the models predicted by various servers and softwares. Default parameters were chosen for running the prediction/modelling and QMean scores were assessed using the site https://swissmodel.expasy.org/qmean/ (Benkert et al., 2009)

| S.No. | Software | Method | QMean4 |
| --- | --- | --- | --- |
| 1 | iTasser | Ab initio | -13.03 |
| 2 | Rosetta | Ab initio | -2.21 |
| 3 | Swissmodel | Modelling | -3.57 |
| 4 | hhPred | Modelling | -5.21 |
| 4 | Schrodinger | Modelling | -7.62 |
| 5 | Raptor | Modelling | -8.20 |
| 6 | Phyr | Modelling | -15.02 |
| 7 | MMM | Modelling | -4.53 |

The ab initio structure predicted by Robetta was energy minimized using the protein preparation wizard of schrodinger and structure quality analysis was performed using the SAVES server (Pontius et al., 1996). The quality of the model was good with 81.18% of the residues with averaged 3D-1D score >= 0.2. The model showed 85.4% of the residues within the most favored regions (red), 12.6% residues within the additionally allowed regions (yellow), 0.7% within the generously allowed regions (beige), and only 1.3% residues of modeled proteins in the disallowed regions (white) (Figure 5a). Thus, the overall stereochemical properties of the model was reliable and was used for further studies. A very basic amino terminal end that is unstructured for more than 30 residues is a characteristic feature of the CP model. Analysis using Protein Disorder prediction System (PrDOS, (Ishida & Kinoshita, 2007), Figure 5b) also suggested that the disorder at the amino-terminus (Figure 4b). Analysis of nucleic acid binding using BindUP server (Paz et al., 2016) (Table 2) also strongly indicated the role of the highly basic amino terminal end towards recognition and binding of cognate genome (Figure 6a). A total of 13 basic residues are present amongst the first 30 amino acids at the amino terminus, mostly featuring lysines. The fold seems to be similar to the β-sandwich fold of the CPs of other plant viruses (Figure 6b) consisting of two β-sheets packed against each other. Sheet 1 is seen to comprise of the three strands A, F and G while sheet 2 comprises of strands B, D. Two short helices are present, α1 between βA and βB and α2 between βE and βF. Strand A is the shortest comprising of 4 residues while strand βF is the longest comprising of 15 residues.

**Table 2:** The nucleic acid binding capacity of the amino terminal 26 residues as predicted by the BindUP server (Paz et al., 2016). POS: Index of amino acid residue, RES: Amino acid residue, S_LBL & S_PRB: SVM prediction of binding label & probability, K_LBL & K_PRB: KLR prediction of binding, label & probability, P_LBL & P_PRB: PLR prediction of binding label & probability, MAJ_CON: Majority consensus, STR_CON: Strict consensus, Labels 0/1 stand for non-binding/binding, respectively, Label NA stands for “label cannot be assigned”.

| POS | RES | S_LBL | S_PRB | K_LBL | K_PRB | P_LBL | P_PRB | MAJ_CON | STR_CON |
| --- | --- | --- | --- | --- | --- | --- | --- | --- | --- |
| 3 | R | 1 | 0.6481 | 1 | 0.6208 | 1 | 0.7480 | 1 | 1 |
| 4 | Y | 1 | 0.6295 | 1 | 0.7573 | 1 | 0.7165 | 1 | 1 |
| 5 | P | 1 | 0.6589 | 0 | 0.6004 | 1 | 0.5896 | 1 | NA |
| 6 | K | 1 | 0.8477 | 1 | 0.8406 | 1 | 0.8695 | 1 | 1 |
| 7 | K | 1 | 0.9378 | 1 | 0.9687 | 1 | 0.8411 | 1 | 1 |
| 8 | S | 1 | 0.8557 | 1 | 0.6821 | 1 | 0.5779 | 1 | 1 |
| 9 | I | 1 | 0.8306 | 1 | 0.9072 | 1 | 0.6931 | 1 | 1 |
| 10 | K | 1 | 0.9576 | 1 | 0.9821 | 1 | 0.8328 | 1 | 1 |
| 11 | K | 1 | 0.8083 | 1 | 0.8845 | 1 | 0.7831 | 1 | 1 |
| 12 | R | 1 | 0.7560 | 1 | 0.7920 | 1 | 0.7163 | 1 | 1 |
| 13 | R | 1 | 0.8760 | 1 | 0.9033 | 1 | 0.7962 | 1 | 1 |
| 14 | V | 1 | 0.7354 | 1 | 0.7259 | 1 | 0.5227 | 1 | 1 |
| 15 | G | 1 | 0.7321 | 1 | 0.6603 | 1 | 0.5968 | 1 | 1 |
| 16 | R | 1 | 0.9340 | 1 | 0.9737 | 1 | 0.8899 | 1 | 1 |
| 17 | R | 1 | 0.9484 | 1 | 0.9439 | 1 | 0.7992 | 1 | 1 |
| 18 | K | 1 | 0.7806 | 1 | 0.6878 | 1 | 0.8730 | 1 | 1 |
| 19 | Y | 1 | 0.8928 | 1 | 0.9688 | 1 | 0.8482 | 1 | 1 |
| 20 | G | 1 | 0.6659 | 1 | 0.5998 | 1 | 0.7547 | 1 | 1 |
| 21 | S | 1 | 0.8290 | 1 | 0.8448 | 1 | 0.8415 | 1 | 1 |
| 22 | K | 1 | 0.7437 | 1 | 0.7864 | 1 | 0.7730 | 1 | 1 |
| 23 | A | 1 | 0.7772 | 1 | 0.6564 | 1 | 0.6591 | 1 | 1 |
| 24 | A | 1 | 0.6096 | 1 | 0.5350 | 1 | 0.5889 | 1 | 1 |
| 25 | T | 1 | 0.6709 | 1 | 0.5969 | 1 | 0.5342 | 1 | 1 |
| 26 | S | 1 | 0.6940 | 1 | 0.6322 | 1 | 0.6782 | 1 | 1 |

**Table 3:**
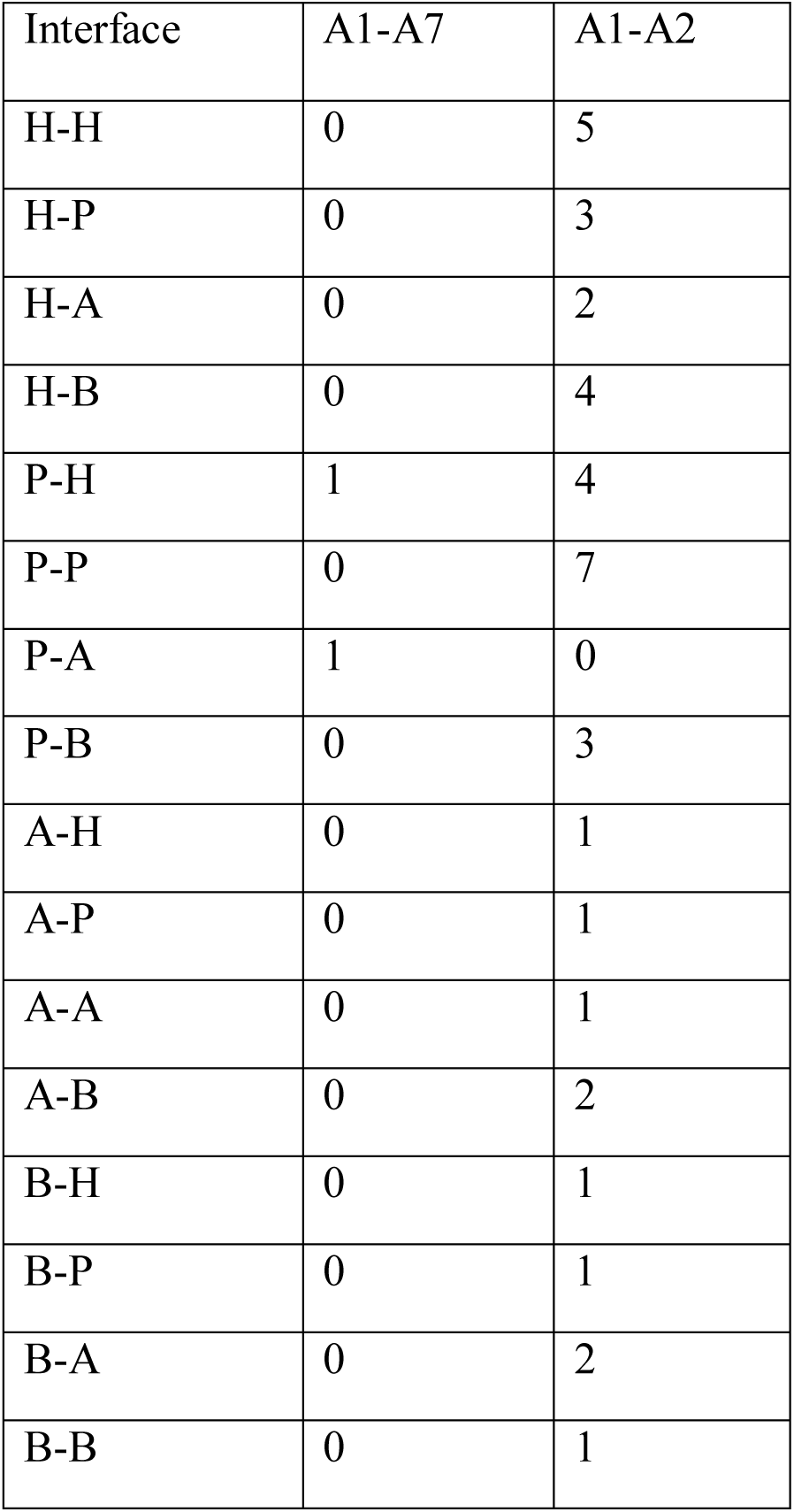
Contacts at the fivefold interface of the capsid as evaluated using the VIPERdb server (Ho et al., 2018). A stands for acidic, B for basic, H for hydrophobic and P for polar.

**Figure 5:**
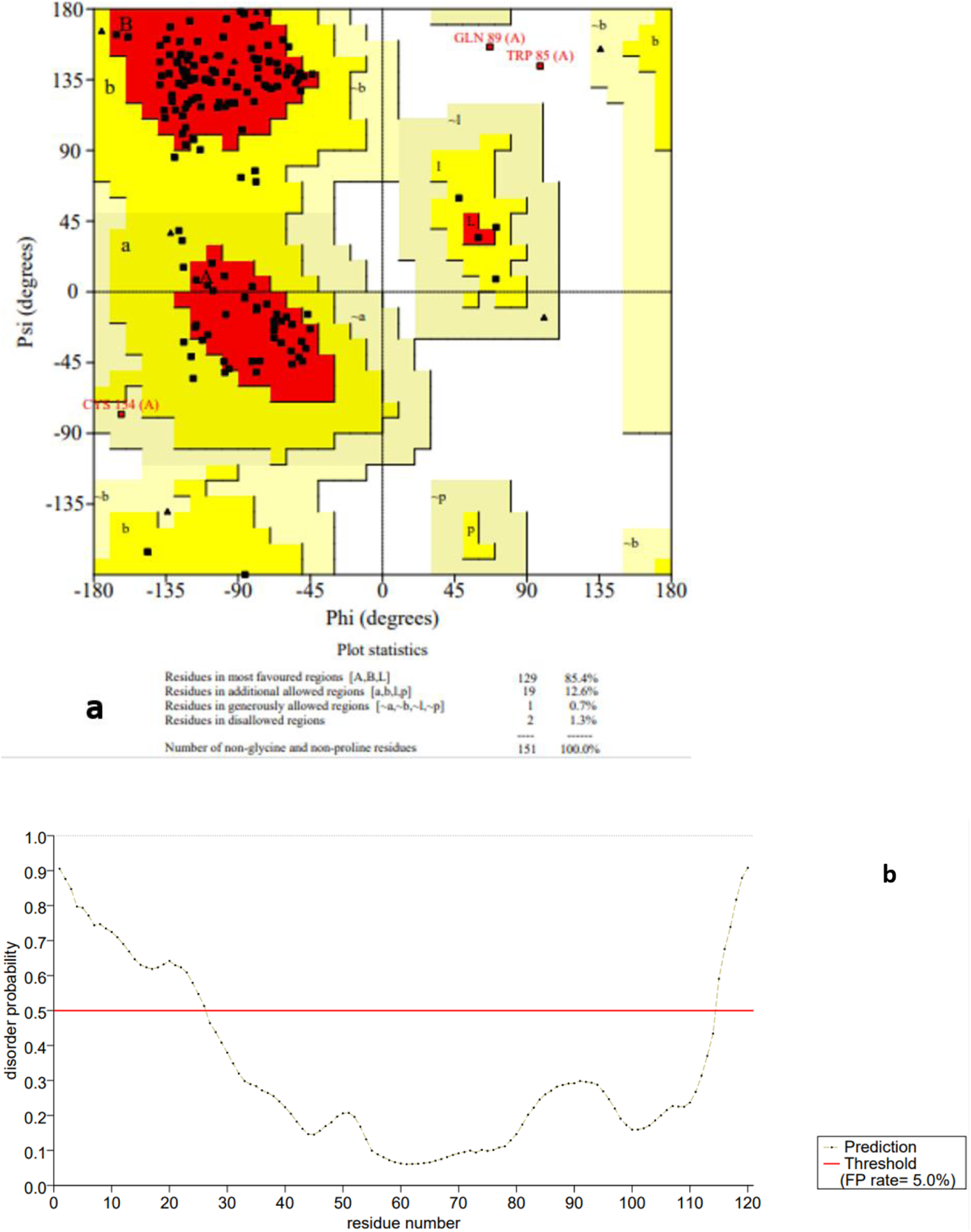
a) Predicted model quality as determined by PROCHECK using the SAVES server (Laskowski et al., 2012; Pontius et al., 1996). More than 82% of the residues are seen in the most favorable region. b) Disorder prediction by PrDOS server that shows a highly disordered amino terminal region (Ishida & Kinoshita, 2007).

**Figure 6:**
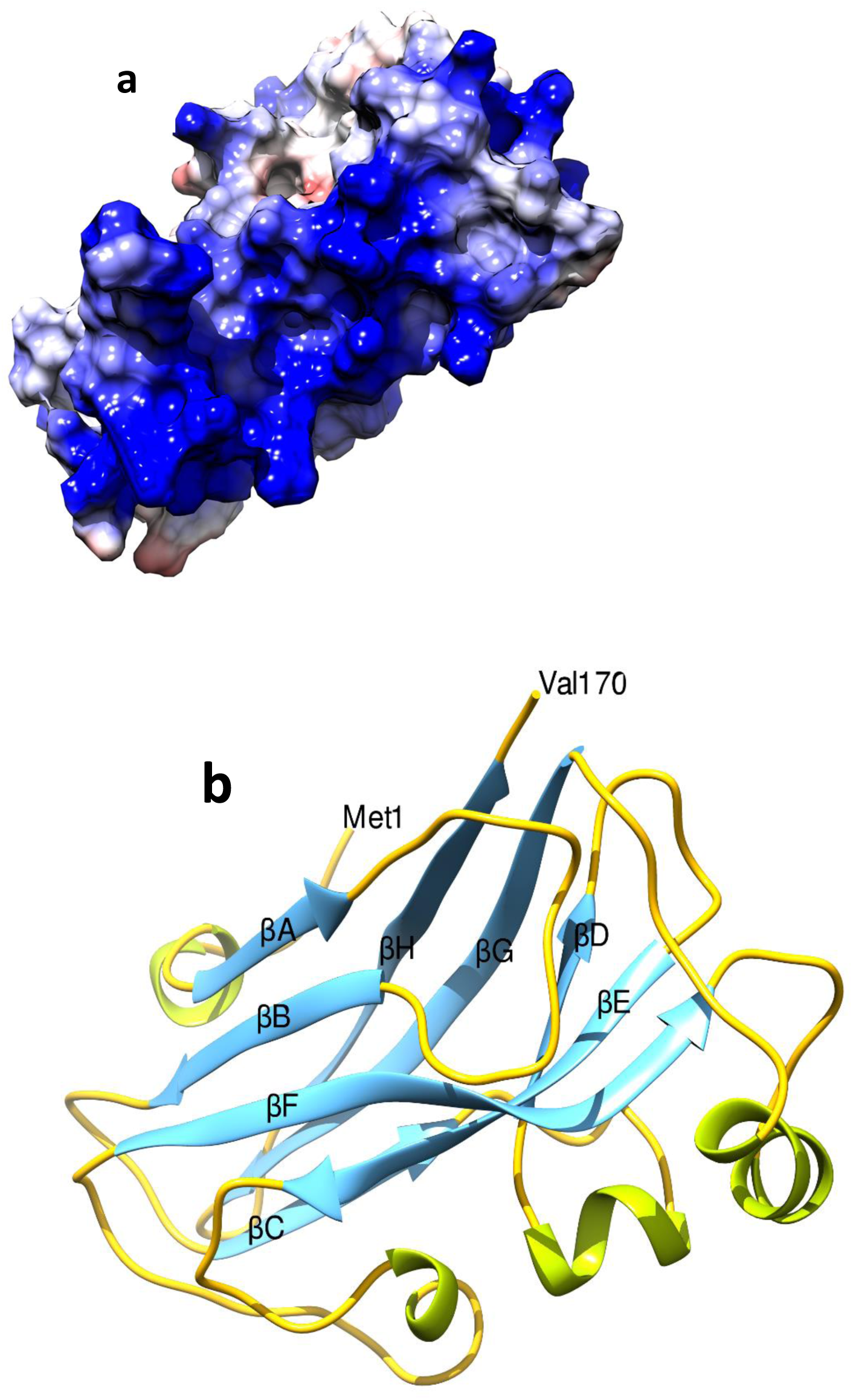
a) Surface representation of the CP showing the highly basic amino terminal region in blue (acidic red and neutral white) that is implicated in binding to the nucleic acid. b) Cartoon representation showing the β-sandwich fold of the CP. The β strands are coloured in blue, α helices in green and loops are in yellow.

### Structure analysis

To study the capsid assembly, the model coordinates of BBTV CP were transformed into PQR system using VIPERdb (Carrillo-Tripp et al., 2009; Ho et al., 2018) in which the icosahedral 2-fold axes are along the P, Q and R directions. The PDB to VIPER transformation matrix is:

0.309 0.809 −0.500

-0.809 0.500 0.309

0.500 0.309 0.809

With a translation of 0.0 0.0 0.0.

The resultant coordinates of the model were used to generate the capsid shell using the 60 icosahedral matrices through the VIPERdb server and the structure was subjected to VIPER analysis (Carrillo-Tripp et al., 2009; Ho et al., 2018). The capsid thus obtained was a closely packed T=1 icosahedron with an average particle radius of 97.0 Å that matches with the diameter of the particles seen under EM (Figure 7a). The model capsid also displays a smooth contour similar to that observed for the natural virus. The outer radius of 97.0 and inner radius being 58.0 Å is consistent with the observed particle diameter in EM studies (Harding et al., 1991; Qazi, 2016). The spherical volumes based on the standard formula, 4/3(Pi*Radius^3) estimated from the average radius was 3,822,995.7 Å^3^. The solvent accessible surface area (SASA) was estimated to be 5,143.7 Å^2^ per subunit of the capsid. The net surface charge was calculated by adding the charges of residues that are surface-exposed (both positive and negative) was estimated to be +900e per virion indicating a highly positive surface. The modelled capsid was analyzed for short contacts and association energies. The contact table (Table 2) indicates a total of 40 unique contacts with seven polar contacts, five hydrophobic and two salt bridges at the A1-A2 interface. The association energy between the A1-A2 interface area is −44Kcal/mol with a buried surface area of 2184.1 Å^2^ and solvation energy of - 20.2 Kcal/mol. Overall the viral capsid resembles a loosely tethered association of twelve pentamers linked by feeble inter-pentameric contacts. This aspect is in agreement with the fact that in the DLS studies one could see a large proportion of pentameric population. The lack of strong inter-pentameric contacts and association is likely the reason that the particles are not readily observable either *in vivo* or *in vitro* conditions. The amino terminal disordered region is seen at the interface lining the pentamers and facing the interior (Figure 6a). It is most likely that nucleic acid mediated interactions play a major role in holding the pentamers together against the otherwise wobbly and basic amino terminal end. Further, BBTV which is primarily transmitted by aphids, lacks the conventional DAG motif in the CP. But instead the EAG motif (residues 118-120) is present and strategically positioned in the highly accessible loop between βF and βG at the pentameric axes in the capsid model (Figure 7b). As previously suggested, EAG could be a potential aphid binding motif and this is conserved in BBTV as well as ABTV. Till date attempts by various international groups to raise monoclonal antibodies against BBTV have been unsuccessful indicating the difficulties in raising antisera for detection of the virus (Mandal, 2010; Qazi, 2016). The predicted capsid structure of BBTV presents valuable insights into why they are inherently unstable and how they contribute towards vector transmission,

**Figure7:**
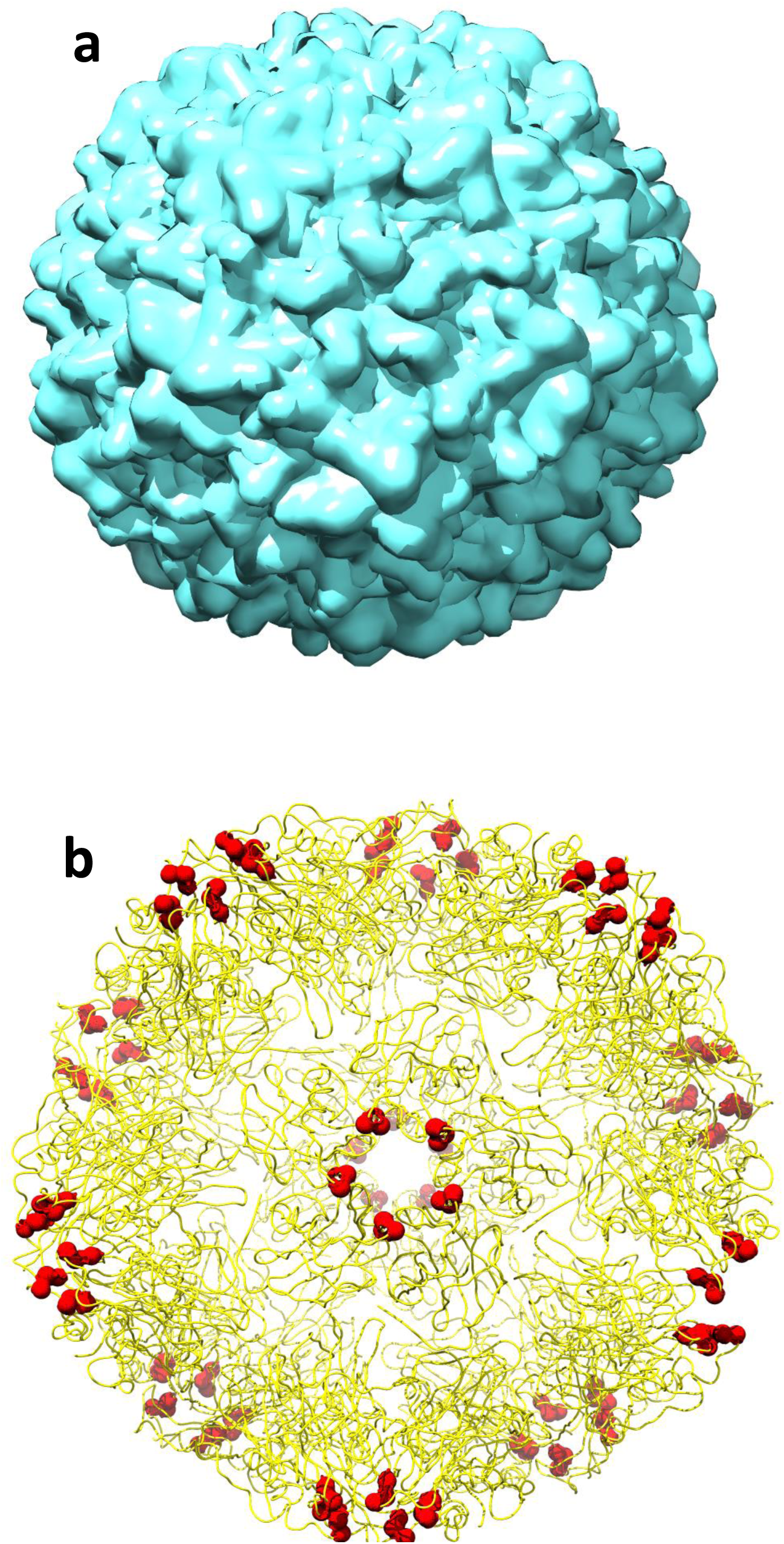
a) Surface representation of the capsid looking down the five fold axis of the icosahedron. b) The aphid binding motif EAG shown as red sphered located at the highly accessible loop between βF and βG at the pentameric axes.

## Conclusion

BBTV is an economically important virus infecting banana plantation around the world. An attempt to understand the structural features of this small virus of 18nm diameter possessing a T=1 shell comprising of 60 subunits of CPs of 19 KDa, the CP gene was cloned into pET28a expression vector and was expressed in E.Coli BL21 DE3 cells. The protein was purified using Ni-NTA ion exchange chromatography. The low yield of the protein hampered crystallographic attempts. Meanwhile, in silico structure determination has been undertaken to predict the structural aspects and derive some idea on possible aphid binding motifs for long term plans to control BBTV infestation.

## Acknowledgement

SV acknowledges the support of SERB for funding the study (ECR/2016/000242).

